# Ultrastructural analysis of the brain endothelium by electron tomography

**DOI:** 10.1101/2024.09.09.612128

**Authors:** Pavel Kotchetkov, Baptiste Lacoste

## Abstract

Transmission electron microscopy (TEM) is a powerful imaging technique, yielding ultrastructural investigation of organic and non-organic samples. Despite its ability to reach nanoscale resolutions, conventional TEM presents a major disadvantage by only acquiring two-dimensional snapshots, thus hindering our volumetric understanding of samples. Electron tomography (ET) overcomes this limitation by offering detailed views of a thin specimen in 3 dimensions (3D). This technique is widely used in biology and has expanded our understanding of mitochondrial structure or synaptic organization. Proper brain functioning is highly reliable on a constant nutritional support through its microvasculature lined by endothelial cells. These unique cells form a selective and protective barrier, known as the blood-brain barrier (BBB), which limits the entrance of blood-borne molecules into the brain. In pathological conditions, the BBB is disrupted, resulting in neuronal damage. Understanding the fine changes underlying BBB disruption requires advanced imaging tools such as ET, to detect the finest changes in endothelial ultrastructure. This manuscript briefly explains how TEM and ET function, and then provides a detailed, didactic method for sample preparation, tomogram generation and 3D segmentation of brain endothelial cells using ET.

## 1 Introduction

### 1.1 Light microscopy vs. transmission electron microscopy

In general terms, a microscope consists of a source of illumination, a lens system that is responsible for formation and magnification of the specimen image, a stage for specimen’s mounting, as well as a device to visualize and store the image^1^. Fundamentally, the absolute purpose of a light microscope (LM) and an electron microscope (EM) is similar, both magnify the image of the specimen. The main distinguishing feature between LM and EM consists in the source of illumination. In LM, the illuminating source is a beam of light (i.e., beam of photons) which can come from either below or above the specimen. The light is diffracted by the specimen, enters the lens which sharply bends light from its original direction and focused. This will generate a true image of the original object. In EM, the illuminating source is a beam of electrons, coming from above the specimen. Electrons entering the electric or magnetic field undergo a deviation similar to the light in LM, however in EM the beam is bent in a continuous process, compared to the sharp refraction of the light in LM.

The major difference between light to electron illumination in microscopy is first and foremost the resolving power. Resolution is the minimum distance between two distinct points, which can be seen separately. It is directly proportional to the wavelength of the illumination source, according to Abbe’s formula. In other words, shorter wavelengths translate into higher resolving power. As an application, visible light with a wavelength of 750 nm gives a resolving power of about 200 nm, while an electron beam at an acceleration voltage of 40 kilovolts (kv) possesses a wavelength of 0.006 nm which results in a resolving power of around 0.5 nm. But since electrons have a much smaller penetration depth than light, EM preparations have to be ultrathin, typically on the nanometers scale^1,2^. The lens systems are very different as well; while LM uses optical lenses, EM uses electromagnetic ones, since electrons are controlled using electric or magnetic fields.

A transmission electron microscope consists of the following components: the illuminating system, the specimen chamber, the image forming system, the image recording system, vacuum system and cooling water circulation. Altogether, these components are assembled in a system, that confer powerful properties to the TEM. Briefly, the illumination system of an electron microscope is composed of the electron gun and condenser lenses that generate and focus electron beam on a specimen, respectively. A set of condenser lenses controls both the intensity and diameter of the beam for high-magnification imaging. The image-forming lens system consists of the objective, intermediate and projector lenses. The objective lens forms the first inverted image and has an influence on the resolution that will be achieved. Further magnification is performed by the intermediate and projector lenses that finally project the image onto a fluorescent screen or a digital camera. The transmission electron microscope works in a vacuum to prevent the interaction of electrons with the medium and to allow clear imaging without air contamination. All these features, in combination, permit elaborate visualization of structural details in anatomy, physiology, and pathology. A conventional TEM can also be upgraded by adding a motorized goniometer. Integrated with specimen chamber, it allows specimen tilting and rotation, which is required for electron tomography^2^.

### 1.2 What is electron tomography?

The term “tomography” originates from the Ancient Greek “*tomos graphos*” for “writing by sections”. The purpose of tomography is to obtain a 3D density distribution of an object, by combining 2D snapshots of this object, acquired from different perspectives. The concept was proposed by Austrian mathematician Johann Radon in 1917; however, the first 3D tomographic reconstruction was performed in 1956 in astronomy. Briefly, tomography consists in capturing 2D images of the object from different projections, and then merging the images to render a detailed 3D structure. Since 1960s, this method began to be actively used in practice in life and material sciences. Further, the application of this method was extended to biological and medical fields, leading to the development of electron tomography (ET) in 1968^3^ and computed tomography (CT) in 1973^4^. Both techniques share the same concept with the difference in the relative motion of the specimen and the source–detector arrangement. ET consists in specimen tilting and a stationary electron beam, and CT scanning consists in X-ray tilting while the patient remains stationary^3,4^.

The application of ET in biology has opened new horizons for the understanding of fine, subcellular structure/function. For instance, ET played a crucial role in structural representation of mitochondrial cristae. Until the 2000’s, the literature used a baffle cristae model, proposed by Palade, where the inner membrane was represented as a continuous complex morphology with cristae, that resulted in large folds, similar to the bellows of an accordion^5^. Since the baffle model was based on 2D single-layer snapshots using conventional TEM, the 3D volumetric information of the mitochondrial cristae remained unclear. By applying ET to mitochondria in 2000, Frey et al. proposed a new cristae model, where narrow tubular openings, called crista junctions, are connected to the intermembrane and intercellular spaces^6^. The ET technique has since then widely expanded our understanding of neuronal structure and neurotransmission. In particular, it provided new insights into the organization of presynaptic vesicles at axon terminals. Harlow et al. applied ET to the frog neuromuscular junctions and demonstrated that active zone material is crucial for synaptic vesicle docking and synaptic channel anchoring^7^. ET methods have also been used to study the organization of postsynaptic density in inhibitory synapses^8,9^ and the arrangement of glutamate receptors at excitatory synapses^9,10^. Overall, ET has become a common tool in neurobiology.

### 1.3 Why focusing on brain endothelial cells?

In the fields of neurobiology and neuropathology it has been applied with a focus on neuronal cells. Yet, to support their crucial role in brain functioning, behaviour and consciousness, neurons are highly reliable on a continuous energy supply and a safe environment, which is provided through regulation of brain homeostasis by blood vessels. Maintenance of brain homeostasis represents a fundamental prerequisite for the optimal functioning of the central nervous system. Brain homeostasis is required for optimal regulation of neurotransmitter levels, oxygen delivery, and pH balance^11-13^. All these factors are directly reliant on cerebrovascular integrity^12,14-17^. Consequently, the cerebral vasculature possesses unique properties, which sets it apart from the peripheral vasculature. In particular, brain endothelial cells are unique in: (1) Absence of fenestrations; (2) Limited paracellular transport via specialized inter-endothelial junctions; (3) Low rate of transcellular transport (transcytosis); and (4) Expression of highly selective influx/efflux transporters^18-20^. Of note, the ATP-dependent nature of many of these transporters underscores a significant energy requirement, which is consistent with the observation that cerebral endothelial cells possess a four-fold greater number of mitochondria compared to peripheral endothelial cells^21^. Altogether, the unique features of brain endothelial cells provide the brain with an impermeable, selective, and protective barrier, known as the blood-brain barrier (BBB). Inter-endothelial junctions are composed of tight (TJs) and adherens (AJs) junctions. TJs are composed of a group of transmembrane proteins that are localized on apical surface of endothelial cells and establish intercellular contact points, thereby limiting the diffusion of ions and molecules across the paracellular pathway. TJs are composed of claudins, occludins and junctional adhesion molecules (JAM’s). Claudin-5 (Cldn-5) is the most critical TJ protein^19,22,23^. All these components play a crucial role in maintaining BBB integrity. Moreover, recent evidence suggests that the BBB plays a crucial role in restricting cellular trafficking, including immune cells. Compared to peripheral tissues, the central nervous system (CNS) has low immune cell surveillance due to the low expression of leukocyte adhesion molecules (LAMs) on CNS endothelial cell. This creates a protected brain environment that is isolated from the immune system, which is referred to as "immunoprivileged"^24-28^. In pathological conditions, such as neurodegenerative disorders, brain tumors, epilepsy, stroke, and inflammation, both para- and trans-cellular transports are affected, leading to the loss of barrier properties^29-32^. Upon loss of BBB integrity, blood-borne pathogens and immune cells can infiltrate the brain parenchyma, leading to neurotoxicity^15-17,33^.

Getting access to the complex ultrastructural features underlying BBB maintenance and disruption is a critical step to improve our understanding of cerebrovascular plasticity, which ultimately will guide therapeutic strategies for neurological disorders. Such advances require imaging techniques that allow for ultrastructural investigation of BBB endothelial cells. TEM is powerful tools to that end^34^, yet conventional TEM is limited to 2D single-layer snapshots of thin specimens, thus lacking 3D insight, and thus restricting our volumetric understanding of the BBB^3,35^. ET is a method which overcomes this limitation.

High quality sample preparation is a crucial prerequisite for 3D reconstruction of the brain endothelium at the ultrastructural level. The following sections describe room temperature ET (‘RT-ET’), as opposed to cryo-ET, thus involving chemical sample preparation of mouse brain endothelium.

## 2 Sample preparation for electron tomography

To preserve the ultrastructure and maintain contrast, the sample should undergo a critical and highly rigorous preparation process. The tissue must be preserved as close as possible to its native state, while using chemical fixation. This process comprises multiple steps, and any error ultimately impacts imaging quality^36^.

### 2.1 Sample fixation

The purpose of this initial step is to halt all metabolic processes and enzymatic reactions for preventing self-destruction of tissue (also known as autolysis). Fixation stabilizes cellular organization and membranes, preventing cellular damage and consequent morphological alterations. There are two common methods of fixation for ET: physical and chemical. The physical method generally involves the cryofixation. The chemical method is the more commonly used and involves chemicals, called “fixatives”.

The choice of right fixative is based on the specimen’s nature as well as on the type of study. Hence, there is no universal fixative for specific specimens. However, the more widely used are paraformaldehyde (PFA) and glutaraldehyde. Both have a strong fixative effect on proteins. They contribute to the formation of a protein gel which results in cross-links between different proteins. The reaction of cross-linking takes place between free amino acid groups.

These fixatives have distinct properties. The penetrating power of PFA, as well as its ability to halt enzymatic activity, are stronger compared to glutaraldehyde. However, the cross-links formed by PFA are less stable, as the process is reversible. Glutaraldehyde is a fixative with slow-penetrating power, but it forms more stable cross-links and increases contrast^2,37^. Given the properties of both fixatives, a mix containing both glutaraldehyde and PFA produces optimal fixation^2,37^. Based on an existing protocol published previously^38^, we found *in vivo* fixation is a most suitable method for mature mice in which transcardiac perfusion can be successful with less challenges (>P30). First, transcardiac perfusion of 50mL of ice-cold 50mM phosphate buffer saline (PBS) is necessary to clear blood vessel lumen from plasma and red blood cells. PBS perfusion is then followed by 225mL of fixative solution containing 4% of PFA (EMS cat. #15713-S) and 2% glutaraldehyde (EMS cat. #16320), diluted in 0.1M phosphate buffer (PB). The chosen concentration of glutaraldehyde can vary from 2% to 3%; however, glutaraldehyde is known to produce tissue shrinkage, thus impacting ultrastructures^39^. We have established 2% glutaraldehyde as an optimal concentration to study the brain endothelium with optimal structural preservation and contrast ratio. To complete structural preservation as well cross-linking, the harvested fixed brains should be incubated in 4% PFA solution in 0.1M PB at 4°C for 30 minutes, followed by two PBS washes before vibratome sectioning (*section 2.2*).

Note that to remove any impurity, it is crucial to filter the fixative solution before beginning the animal perfusion. More importantly, to prevent the endothelial damage from perfusion pressure, perfusion flow rate should not exceed 5mL/minute.

Chemical fixation for ET can also be performed by immersion without perfusing the animal, and this is recommended for younger animals in which transcardiac perfusion is more challenging (<P21). This method consists of harvesting the brain and immersing it in a fixative solution containing 4% of PFA (EMS cat. #15713-S), 2-3% of glutaraldehyde (EMS cat. #16320) in 0.1M sodium cacodylate buffer pH 7.4 (EMS cat. #11653). To minimize shrinkage, the optimal concentration of glutaraldehyde should not exceed 3%. For small brain parts (e.g., micro-dissected hippocampus, cortex), samples should be immersed in fixative solution for 30 minutes at room temperature. For a whole brain, the immersion should be prolonged to 1 hour. After this first immersion, the brain should be transferred into a fresh fixative solution, prepared with the same recipe, and immersed for 5 hours, followed by rinsing in a cacodylate buffer pH 7.4 overnight at 4°C. Following this overnight rinsing, the brain sample should undergo a last brief rinsing in cacodylate buffer pH 7.4 before performing vibratome sectioning.

Despite the high reactivity of glutaraldehyde and PFA with proteins, both have a selectivity limitation for lipids. Therefore, post fixation is the next step for ET sample preparation, which consists in the contrast enhancement of membranes. However, to enhance penetration of fixatives, the brains should be thin sliced with a vibratome prior to post-fixation.

### 2.2 Vibratome sectioning

Fixed brains should be sectioned free-floating coronally into serial, 50μm-thick sections using a vibratome (e.g., Leica VT1000S). To obtain optimal vibratome parameters for brain sectioning, we set up the frequency and speed at mode “10” and “3”, respectively. 50μm-brain sections can either be stored for long period at -20°C in an anti-freeze solution (30% glycerol, 30% ethylene glycol, 40% of PBS) or used immediately for processing (post-fixation, *section 2.3*). For optimal preservation of the ultrastructure, it is highly recommended to perform vibratome sectioning the same day after the last step of sample fixation (*section 2.1*).

### 2.3 Post-fixation

Osmium tetroxide (OsO_4_) is commonly used as a secondary fixative that interacts with lipids without altering cytoplasmic and mitochondrial morphology. Given its destructive properties on proteins, OsO_4_ cannot be used as a primary fixative^2,36,40^.

Based on existing reports^41-45^, we found that using the osmium-tetracarboxyhydrazide-osmium (OTO) method is the optimal protocol for ET, as it minimizes artifact formation and does not require additional contrast enhancements using heavy metal salts. This method involves double staining with OsO_4_ using tetracarboxyhydrazide (TCH) as a bridge between two treatments with osmium^45^. For the OTO method, described below, samples are first treated with a mix of OsO_4_ and potassium ferrocyanide (K_4_Fe(CN)_6_). Developed by Morris Karnovsky, the reaction between OsO_4_ and K_4_Fe(CN)_6_ results in reduced OsO_2_, thus preventing the formation of precipitates through interaction with lipids. The second treatment with OsO_4_ consists of contrast enhancement through the binding of more osmium using TCH as a bridge between OsO_2_ and OsO_4_^44,45^.

First, stored tissue sections are extracted from the anti-freeze solution and washed in PBS 1M 5 times for 3 min^38^. During washing, two solutions are prepared. The first solution consists of mixing a solution containing 3% K_4_Fe(CN)_6_ (Bioshop, cat. #PFC232.250) diluted in 0.1M PB with an equal volume of 4% aqueous OsO_4_ (EMS, cat. #1950) to obtain reduced osmium dioxide OsO_2_. Tissue sections are then incubated in this mix for 1 hour at room temperature, protected from light (e.g., covered in aluminum foil). The second solution contains 0.1 g of TCH (EMS, cat. #21900) dissolved in 10mL of Milli-Q water and incubated in an oven at 60°C for 1 h (while the samples are incubated in the ferrocyanide-osmium solution). Importantly, for the homogenous dissolution of TCH powder in Milli-Q water, it is recommended to gently agitate the warming TCH solution every 10 min. At the end of the oven incubation, the homogenous TCH solution is filtered through a 0.22 μm Millipore syringe filter before use. After incubation in the reduced osmium dioxide OsO_2_ solution, the samples are washed in Milli-Q water 5 times for 3 min each. After washing in Milli-Q water, tissue sections are incubated in the filtered TCH solution for 20 min at room temperature, also protected from light. At the end of incubation in TCH solution, five washes in Milli-Q water (3 min each) are performed before the next step. Thereafter, samples are incubated for 30 min at room temperature in the dark in a solution containing 2% OsO_4_ diluted in Milli-Q water (i.e., second exposure to Osmium). As mentioned above, this double incubation in osmium solution enhances the contrast by bridging two molecules of osmium. The final post-fixation step consists of another series of washes in Milli-Q water (five times, 3 min each) after the incubation in 2% OsO_4_. Following fixation, the samples are dehydrated.

### 2.3 Dehydration

Since all biological samples contain water, this leads to two major challenges. The first challenge consists in the difficulty to produce ultrathin sections. The second challenge is the incompatibility of loading aqueous specimen in the high-vacuum column of an electron microscope. For these reasons, water in samples should be first replaced by organic solvents via dehydration^40,46^. This step is performed by incubating tissue sections in a graded series of ice-cold ethanol, in the following order: 30%-50%-70%-80%-90%-100%-100%-100%. Each treatment must last for 2 minutes. The final step consists in a total water removal by replacing the last (3^rd^) solution of 100% (anhydrous) ethanol with a propylene oxide (Sigma, cat. #110205-1L) and incubating samples for 5 minutes. Importantly, propylene oxide is a plastic dissolvent, therefore it is important transfer the brain slices into glass vials (EMS, cat. #72632) and use a Hamilton glass syringe at the time of propylene oxide collection. The replacement of anhydrous ethanol with propylene oxide solution also is a prerequisite for optimal resin embedding, as propylene oxide also serves as resin dissolvent.

### 2.4 Resin embedding of samples

The purpose of embedding is to harden the specimen to facilitate ultra-thin sectioning. It consists of the replacement of propylene oxide by a resin monomer which, once polymerized, generates solid blocs of embedded tissue. This contributes to optimal sectioning properties, with a minimal tissue compression. In addition, embedding in resin ensures tissue stability under the electron beam and consequently prevents specimen damage^2,40,47^.

For this step, a mix of four Durcupan resin components (EMS, cat. #14040) is prepared in the following order: 20g of component A, 20g of component B, 0.6g of component C and 0.4g of component C. The mix is slowly and gently mixed without creating bubbles, and then transferred to aluminum dishes (Fisher, cat. #08-132-100). At the end of the dehydration step (i.e., after propylene oxide), tissue sections are transferred to the aluminum dished containing the prepared resin, sunk in resin and incubated overnight at room temperature. Following this overnight incubation, samples are warmed 5 min at 55°C (to liquify the resin), placed between two Aclar sheets (EMS, cat. # 50426-25) embedded in a thin layer of liquid resin, and transferred into an oven for 72 hours at 55°C for resin polymerization.

### 2.5 Ultra-thin sectioning

Due to the electron beam limited penetration, the resolution of EM image is highly dependent on the thickness of sections. The thicker the section, the lower the contrast. An ultramicrotome is the tool used to perform ultrasectionning^2,47-49^. Prior to ultra-thin sectionning, cortical regions are isolated from the resin-embedded brain slices and thoroughly cut onto 2mm^2^ pieces using a scalpel. Subsequently, these pieces are mounted on resin blocks using cyanoacrylate adhesive (Gorilla super glue®).

For ET specifically, ribbons of 140nm-thick ultra-thin sections (e.g., using a Leica UC6 ultramicrotome) are and collected on 100-mesh copper grids. We established that 140nm is an optimal thickness for ultra-thin sections in order to reconstruct the brain endothelium using ET. It can be extended up to 150nm to enlarge the 3D volume of the tomogram; however, we experienced that it makes it more challenging and time consuming to find a region of interest.

Overall, every step mentioned above is crucial for obtaining a sample with a high contrast, high resolution and less artifacts. Any error induced during sample preparation can impact the quality of the sample and subsequently tomogram quality.

## 3 Electron tomography

ET comprises the following main steps: (1) Acquisition of the data by tilting the specimen; (2) Processing of the tilt data by aligning a stack of acquired 2D images and assigning X/Y/Z rotation angles (also known as Euler angles); (3) Computational 3D reconstruction and visualization; and (4) Segmentation and interpretation of the tomogram^50,51^.

### 3.1 Data acquisition by specimen tilting

Acquisition represents a fundamental step for ET as it consists of recording 2D images at various perspectives using a goniometer, by rotating the specimen around the tilt axis^52^. Understanding of key parameters for the acquisition step, such as the accelerating voltage, electron dose, choice of tilt range, tilt increment and tilt scheme will define the quality of the final tomogram, and subsequent segmentation. The balance between these parameters will provide a final tomogram with optimal resolution and contrast and minimized artifacts^53^.

The first parameter to consider optimizing RT-ET is the accelerating voltage, as it impacts both resolution and the contrast. In fact, accelerating voltage and contrast are inversely proportional. By increasing accelerating voltage, electrons gain more energy and are transmitted faster through the sample, leading to the reduced scattering and interaction with the sample, thus resulting in lower contrast. Low accelerating voltage will cause more scattering and interaction with the specimen, leading to the higher contrast. However, high interaction between electrons and the specimen will induce important specimen heating and subsequent molecular motion, which will cause specimen distortion and ultimately destruction. Conversely, resolution is directly proportional to the accelerating voltage as the latter facilitates electron penetration through the specimen, thus minimizing electron scattering and diffraction within the sample. This concept is also applicable for thick specimen, suggesting that thicker specimen require a higher accelerating voltage to collect high-resolution data^54,55^. Accelerating voltage is also closely related to electron dose, another parameter that refers to the amount of electron exposure of the specimen during imaging. The higher the electron dose, the higher the contrast and resolution. However, similar to accelerating voltage, electron dose also negatively impacts the specimen due to heating, causing molecular motion and sample destruction^53,55^. Of note, in the examples discussed hereafter (see **Figure 1**), we used our 120kv JEOL JEM1400-Flash equipped with a bottom-mount Gatan OneView 4k digital camera and a goniometer capable of ±70° tilt angle range. All our tomograms are generated at an acceleration voltage of 120 kv (with a LaB6 filament).

**FIGURE 1.**
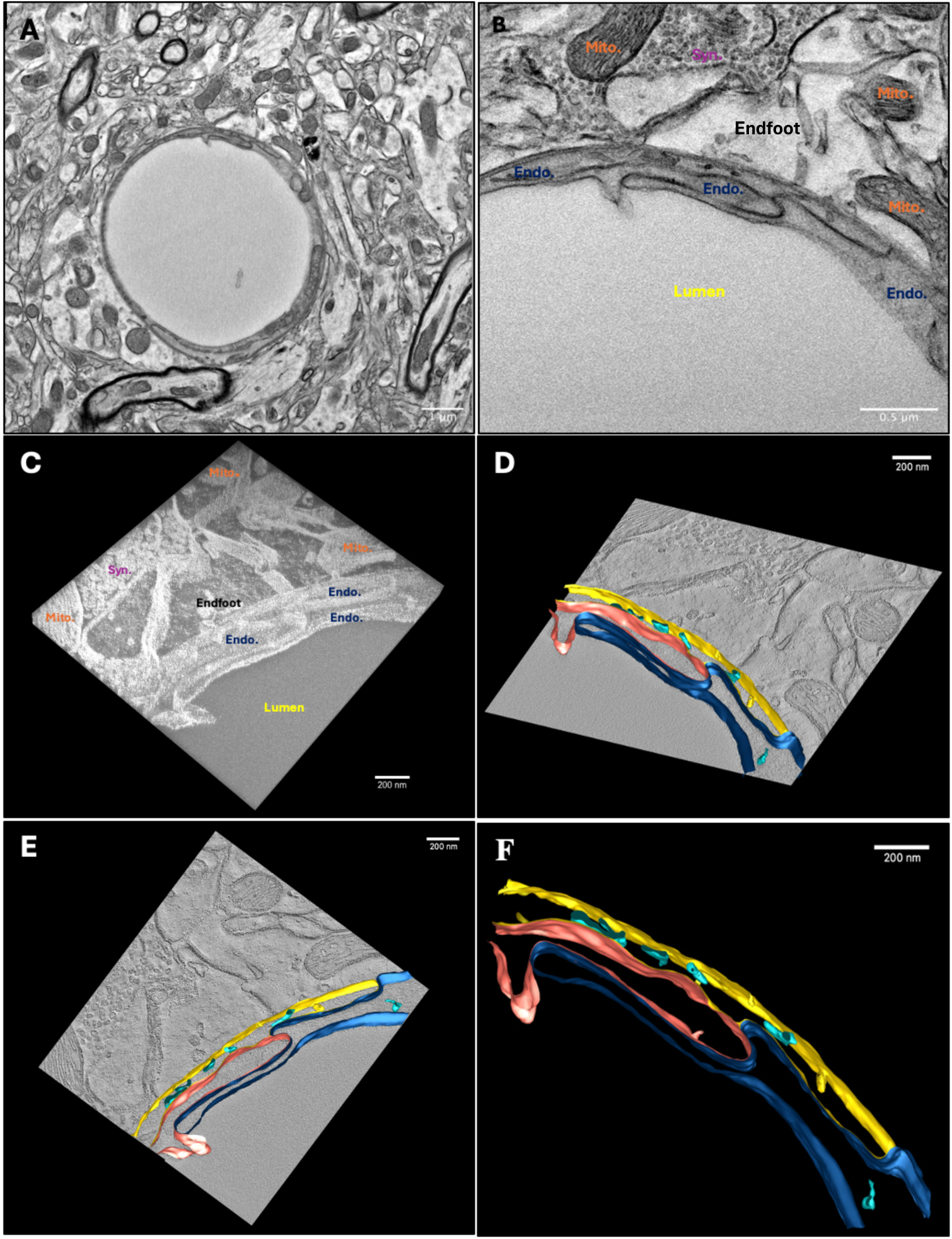
Ultrastructural 3D reconstruction of the brain endothelium using ET. A) Conventional TEM micrograph of a vascular profile, acquired at low magnification from a 140nm-thick section. B) Conventional TEM micrograph acquired at medium magnification, demonstrating three murine cerebral endothelial cells tightly sealed by inter-endothelial junctions. Endo.: Endothelial cell, Endfoot: Astrocytic endfoot, Syn.: Synapse, Mito.: Mitochondria. C) 2D snapshot of reconstructed 3D volume of murine cerebral inter-endothelial junctions. D) and E) 3D-segmented model of murine endothelial components. The tomogram planes are superposed in between the segmented objects. In light blue colour: endoplasmic reticulum. F) 3D-segmented model of murine endothelial components without superposition of tomogram planes.

Another important parameter impacting contrast, resolution, and faithfulness of the tomogram is the tilt angle range and the angular increment selected. The tilt angle range refers to the angle series at which the specimen is tilted during acquisition. Generally, this parameter does not exceed ±60° rotation; but exceptionally it could be extended up to ±80°. Tilt angle range plays a crucial role in specimen coverage as it determines the faithful representation of the object on the final tomogram. Consequently, a wider tilt angle range provides a better specimen coverage and a more complete volumetric representation of the object. Even though a wider tilt angle range is beneficial for providing faithful specimen representation, it also implies longer specimen exposure to the electron beam, which might damage the specimen^50,56,57^. Additionally, a wider tilt angle range can become challenging for thicker specimens. The more the tilt angle increases, the more the projected thickness for electron beam increases upon tilting. Based on these facts, for a 70-nm thick specimen, the projected thickness will represent approximately 140nm at 60°^58^. Furthermore, due to the shadowing originating from the specimen holder, and due to the restricted space in the specimen chamber, the tilt angle range cannot be extended to a complete ±90° rotation, thus making several specimen regions inaccessible to electrons. This phenomenon is referred to as “missing wedge effect”, which is responsible for artifacts such as structural elongation and distortion^53,55,59^. This phenomenon usually occurs while performing a single-axis tilting. This method involves the specimen tilting only around one axis and is commonly used for shorter data acquisition times, as well as less adjustments and processing. Conversely, dual-axis tilting involves the specimen tilting around two perpendicular axes. This method minimizes the “missing wedge effect”, however it requires more time for data acquisition, as well more adjustments and tilt data processing to generate a tomogram.

The tilt angle range is not the only parameter that impacts the completeness of the tomogram; the angle increment is another important parameter to set up before acquiring data. While tilt angle range represents the interval of angles that the specimen is tilted at, the angle increment refers to the angle that the specimen is tilted at each step of image recording. The smaller the angle increment is, the more detailed a volumetric object recording is, thus providing a better volume completeness with minimized artifacts. It also improves data processing, ensuring a higher tomogram resolution. Typically, the angle increment should vary between 1° and 2° to obtain a high-quality tomogram^55,60^.

Both, tilt angle range and increment impact optimal data collection. However, the data can be collected in different order and angles, which is referred to as tilt scheme. There are three different tilt schemes: continuous, bi-directional or dose-symmetric. Continuous tilt scheme consists of starting by collecting data from one extreme tilt angle and terminate the tilting on the opposite extreme angle. This method is not optimal for the large tilt angle ranges, as the specimen gradually degrades by reaching the opposite extreme angle. Additionally, at 0° the projected thickness is minimal, resulting in better contrast and resolution. Therefore, continuous tilt scheme is less effective because of specimen damage at the point when the projected thickness is the most optimal. Bi-directional tilt scheme consists of beginning data collection at 0° and proceeding towards one extreme angle, then returning to 0° and resuming data collection towards the opposite extreme angle. One of the advantages of this tilt scheme is to collect tilt data at minimal projected thickness with a low electron dose, ensuring better resolution. However, this scheme is not optimized for a homogenous electron dose accumulation. Upon return to 0°, the second half of the tilt range has already accumulated electron dose and may cause further specimen damage, thus making subsequent tilt data processing more challenging. Conversely to bi-directional tilt scheme, a dose-symmetric tilt scheme is characterized by tilting a specimen following a pendulum-like motion until the extreme angle. This method ensures a homogenous electron dose acquisition; however, there is a risk of a long sample stabilization after tilting, which can lead to a potential loss of the region of interest during tilting^53,55^.

Altogether, these parameters should always be considered before starting tilt series acquisition, as they will impact the subsequent steps of 3D reconstruction and segmentation.

### 3.2 Processing of the tilt data

Upon termination of the tilting session, the acquired raw tilt data represents an image stack with potential artifacts, such as image shifts along the X/Y/Z axes and magnification changes. These artifacts are due to the automated acquisition and can be corrected for any X/Y/Z shifts through alignment, which can be performed using different methods: fiducial-marker alignment or cross-correlation alignment.

Fiducial-marker alignment involves specimen treatment with colloidal gold nanoparticles. These nanoparticles are tracked and used as a reference to determine Euler angles and positions of the object through projection equations. This step is crucial for geometric parameters that define the spatial positioning of the object of interest according to the tilt axis. Despite a precise alignment, the electron-dense gold nanoparticles create artifacts that can overlap biological ultrastructures^55^.

The alignment of collected tilt data can also be performed using an automatic fiducial-less method through the cross-correlation of common motifs of the specimen between projections. In this case, tilt images are aligned based on the reference image, typically acquired at 0°. During the alignment step, image shifts and distortions may occur, causing artifacts. Therefore, the aligned data is subsequently refined through iterative area matching and coordinate correction. This method ensures a highly accurate automatic alignment and minimizes manual intervention and artifacts. However, ensuring optimal contrast and resolution of the reference is crucial, as it might interfere with subsequent 3D reconstruction. Hence, high-quality specimen preparation is essential for identifying common motifs and improving the accuracy of the alignment^55,61^.

### 3.3 Computational 3D reconstruction

Once the tilt series is aligned and refined, each 2D image in the tilt series should be computationally reconstructed into a complete 3D volume of the object. The 3D reconstruction is performed using back-project algorithms. The concept of back projection involves reconstructing the positions of 2D images by following the path of the electron beam that captured those images originally. However, when translating 2D images into a 3D volume through back projection, uneven processing can occur, which results in visual distortions such as pixel smudging or blurriness particularly noticeable, around the edges of structures. Back projection algorithms are designed to enhance the accuracy of 3D reconstruction by reducing these distortions with various techniques. One of the algorithms used includes the weighted back projection method (WBP), along with the simultaneous iterative reconstruction technique (SIRT) and simultaneous algebraic reconstruction technique (SART)^55,62^.

The WBP technique involves filtering out distortions by modifying the influence of each projection used in the process. Following the acquisition of tilt series images, the projections taken near the objects center are often gathered frequently and hold a greater amount of information (pixels) compared to those collected from the object peripheries. This situation is particularly noticeable when capturing projections at angles within the tilt angle spectrum. Consequently, sections of the item that were gathered frequently would have a greater impact on the 3D reconstruction process, resulting in biased and inaccurate portrayal of the item. The WBP algorithm gives importance to edge pixels compared to center pixels ensuring a consistent portrayal of the object. Although WBP is widely utilized due to its simplicity and quick processing speed, it is heavily reliant on noise levels that could potentially impact the clarity of the generated 3D volume^55,58,62,63^.

The SIRT and SART methods are based on a similar concept: the creation of an estimate of the tomogram through iterative virtual reprojections and calculating the difference between each pixel of virtual reprojections and experimental 2D images. Subsequently, the algorithm adjusts the tomogram by distributing the pixel differences back to the 3D volume through reconstruction. While the SIRT algorithm adjusts the entire tomogram simultaneously, the SART algorithm adjusts each angle successively^55,59,62^.

### 3.4 Segmentation and interpretation of the tomogram

Tomogram segmentation consists in identifying and delineating different structures of interest within a reconstructed 3D volume. Segmentation accuracy is crucial for achieving faithful 3D representation of the specimen. However, important challenges significantly impede the segmentation process: (1) The inherent morphological complexity of specimen makes it challenging to delineate the structures; (2) The specimen distortion due to the electron beam exposure could cause shifts throughout the entire raw tomogram, leading to imprecise 3D reconstruction; (3) The “missing wedge effect”, causing the absence of information in tilt data, makes the segmentation process challenging and time-consuming. It can be performed using different software such as IMOD^64^, Dragonfly^65^, Microscopy Image Browser (MIB)^66^, TomoSegMem^67^, Bsoft^68^. In addition, several computational methods have been developed to facilitate this task^69^. However, the challenges mentioned above still persist, thus making manual method as the optimal yet tedious method of segmentation of brain endothelial cells.

Interpretation of the tomogram is a subjective task, that depends on many factors. The major factor consists in the quality of sample preparation and tomogram generation, ensuring the minimum of artifacts, inducing bias. Another essential key of accurate interpretation is a correct identification of ultrastructure, guided by user knowledge. Here, we present a 3D rendered model of three adjacent brain endothelial cells tightly sealed by inter-endothelial junctions, in a healthy adult (P50) male C57BL/6 mouse (**Figure 1**). The junctional strands appear uniform, consistent and evenly distributed, without structural abnormalities such as gaps or tortuosities that would be observed in pathological conditions such as stroke^34^. In addition, these cerebral endothelial cells exhibit cytoplasmic and membrane bounded transcytotic vesicles with normal morphology, and highly abundant endoplasmic reticulum (highlighted in light blue colour). Altogether, these criteria are commonly used for the structural investigation of brain endothelial cells.

## 5 Tomogram generation, segmentation and 3D rendering

Below we provide a protocol comprising tilt data acquisition, tilt data alignment, tomogram generation, and 3D segmentation for room-temperature ET.

### 5.1 Tilt data acquisition

Here, we provide a protocol for ET of mouse brain endothelium using the SerialEM software, developed by David Mastronarde^70,71^. Prior to tilt series acquisition, several conditions should be verified. First, it is essential to ensure that the integrity of ultra-thin sections is consistent and covers the entire square of the grid at minimal magnification (x200). The presence of folds or fractures within ultra-thin sections will cause specimen shift and even complete disruption, thus making it impossible to continue the experiment. Furthermore, brain microvascular profiles must be identified and ensure that they do not exhibit any morphological deformation. It is also essential to optimize the focal plane and align the beam to the center of the column, as this will impact contrast and resolution during the procedure.

Once ultra-thin section and microvascular integrity is verified, the angle tilt range should be validated so the field of view does not overlap with grid bars. SerialEM possesses a tilt control panel, allowing precise navigation to the desired angle within the angle tilt range. The user should navigate to the extreme angle by clicking “To” and set +60°. After clicking “OK”, the goniometer will rotate to the chosen angle. This procedure should be executed for both extreme angles (±60°). Of note, it is acceptable if the overlap occurs closer to the extreme angles of the tilt range, between ±55° and ±60°. This overlap can be removed from tilt data using a crop option during the 3D reconstruction step, as discussed in section 5.3. In our example, tilt data acquisition was performed at x25000; however, the magnification should be adjusted according to specimen morphological specificities. Before initiating the tilt series, it is important to calibrate the eucentric height so that the region of interest remains in the center of the field of view and in focus during the tilting process^72^. This step is achieved first by calibrating a rough eucentricity to get the region close to the eucentric height (5-10μm). This step can be performed by navigating to the menu bar and pressing “tasks → eucentricity → rough eucentricity”. After the rough eucentricity is calibrated, a fine calibration should be performed by selecting “eucentricity → refine and realign” (**Figure 2A**).

**FIGURE 2.**
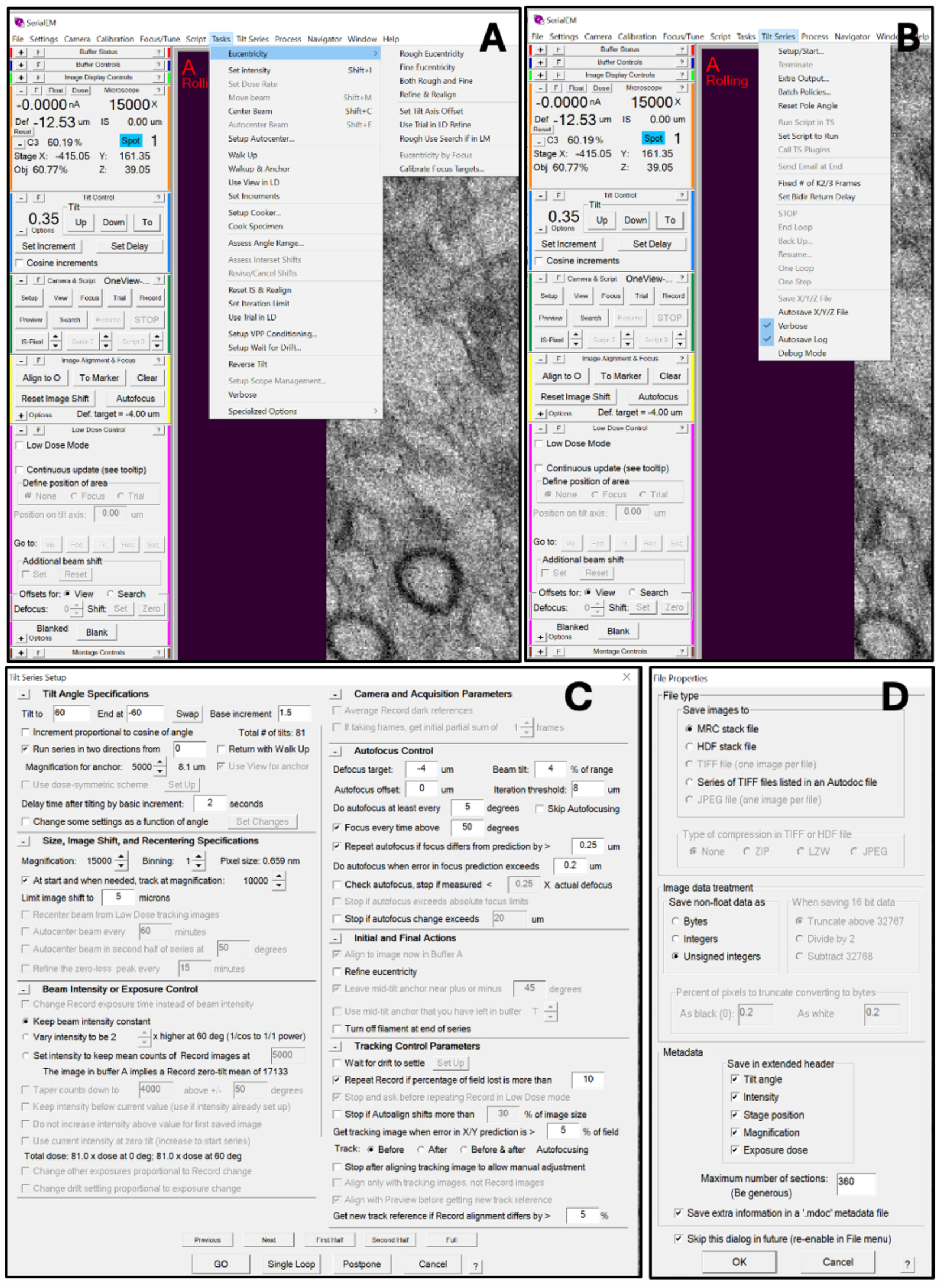
Tilt data acquisition using SerialEM software. A) Calibration of rough and fine eucentricity. B) Start of the tilt series acquisition. C) Tilt angle specifications window, offering to set tilt range and increment angle. D) File properties window.

Once the optimal eucentric height is set up, tilt series acquisition can be started by selecting the “tilt series → setup/start” command (**Figure 2B**). A new window displaying tilting parameters will appear (**Figure 2C**). By selecting the “GO” command, the software will prompt to specify the name and the directory for the file containing the from now on acquired tilt data and save as an *.st* file. Before starting the tilting process, a new window will appear and offer to select file properties. The user should check the following parameters: “MRC stack file”, “Unsigned integers”, “Tilt data”, “Intensity”, “Stage position”, “Magnification” and “Exposure dose”. Assign the “Maximum number of sections” parameter to “360” and select “Save extra information in a ‘.mdoc’ metadata file” and start tilting series by pressing “OK” (**Figure 2D**). The acquisition process will first begin by navigating to the region of interest at a low magnification (x6000). After that, the magnification will increase to the level set by user, thus shifting the region of interest off-axis. This process ensures the beam is transmitted over the large area of the specimen, thus minimizing heating of the region and subsequent damage during recording. The beam is then “blanked” while focusing, and after setting the defocus, the snapshot is acquired. This process is repeated automatically at every tilt angle until completing the tilt series acquisition. After being acquired, these projections undergo alignment and reconstructed through various back projection algorithms.

### 5.2 Tilt data alignment

To perform alignment of the acquired tilt data, we use the Etomo (IMOD) software, developed by David Mastronarde^73^ and following these steps: pre-processing, coarse alignment, tomogram positioning, and creation of final aligned stack.

After launching Etomo, the user should select command “Build tomogram” from the front page and select saved *.st* file containing tilt data in the “Raw Image Stack” window. After selecting “Scan header”, the software will determine pixel size and image rotation. Even though we used a fiducial-less method of alignment, the software requires to set the fiducial diameter: “10” is a number used by default. After selecting the axis type (“single axis” in our experiment), the alignment step can be started by selecting “Create Com Scripts” (**Figure 3A**). A new window will appear showing different steps of tomogram generation. Select “Pre-processing → Create Fixed Stack → Use Fixed Stack” to apply and save the modifications (**Figure 3B**). After clicking “Done”, the software will proceed to Coarse Alignment step. Select “Calculate Cross-Correlation → Generate Coarse Aligned Stack” and check “Coarse alignment only”. Press “Done” to proceed to tomogram positioning and generation (**Figure 3C**).

**FIGURE 3.**
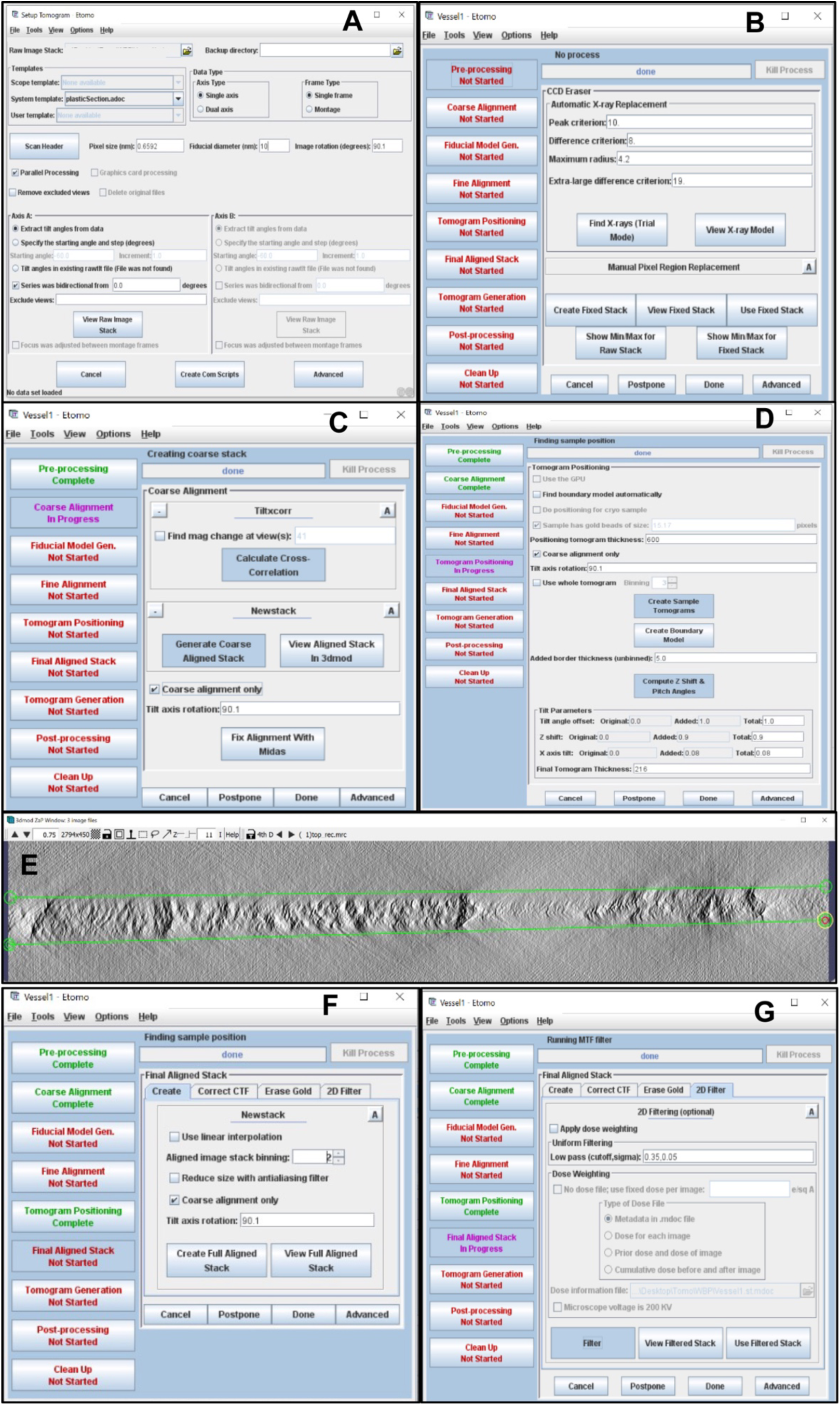
Tilt data alignment using Etomo software. A) Start window of Etomo software. B) Creation of fixed stack. C) Creation of coarse stack. D) Tomogram positioning and creation of boundary model. E) Created boundary model of the top of the tomogram. F) Creation of full aligned stack. G) Optional application of the 2D filter.

Set the “Positioning tomogram thickness” to “600” to generate a reconstruction with visible borders, then select “create Sample Tomograms → Create Boundary Model” (**Figure 3D**). A new window will display the reconstructed surfaces of the tomogram top, middle and bottom. Delineate the tomogram boundaries for all levels: top, middle and bottom, by pressing on mouse middle button (**Figure 3E**). Once the boundaries are set up at all levels, save the model file and press on “Compute Z Shift & Pitch Angles” so the algorithm can calculate the tilt parameters and define the final tomogram thickness. After pressing “Done”, the software will create the final aligned stack. To reduce the time of tomogram generation and the volume of the final tomogram, assign “Aligned image stack binning” to “2” for faster data processing and data size reduction. Afterward, press “Create Full Aligned Stack” to terminate the alignment step (**Figure 3F**). Optionally, if the image displays excessive noise, a 2D filter can be applied by selecting “Filter” and “Use Filtered Stack” (**Figure 3G**).

### 5.3 Tomogram generation

Following tilt series and tilt data alignment, the final step consists of tomogram generation. This procedure can be performed through back projection or SIRT algorithms. For our experiment, we generated tomogram using weighted back projection algorithm. The user should set Central Processing Unit number (#CPUs) to a maximal value so the software utilizes the full processor capacity for faster processing and press on “Generate Tomogram” (**Figure 4A**). Once the process is done, the software will proceed to post-processing of the tomogram, allowing users to trim volume and exclude blurry tilt angles. After navigating to the post-processing step, select “3dmod Full Volume” to view the generated tomogram. A new window containing 3D reconstructed volume will appear with a menubar on the top (**Figure 4B**). Verify whether the 3D volume contains blurry limits by scrolling through Z-axis range towards the extremities. By selecting “rubber band icon”, situated on the left from Z-axis range, two icons “LO” and “HI” will appear. These allow to select low and high Z values for the reconstructed volume and exclude blurry images. To do that, scroll Z-axis to the left until the image becomes blurry and press “LO” to set the low Z limit and repeat the process to define the high Z limit by pressing “HI”. Once the Z limits are set appropriately, the reconstructed 3D volume can be cropped by drawing a region of interest with mouse left button.

**FIGURE 4.**
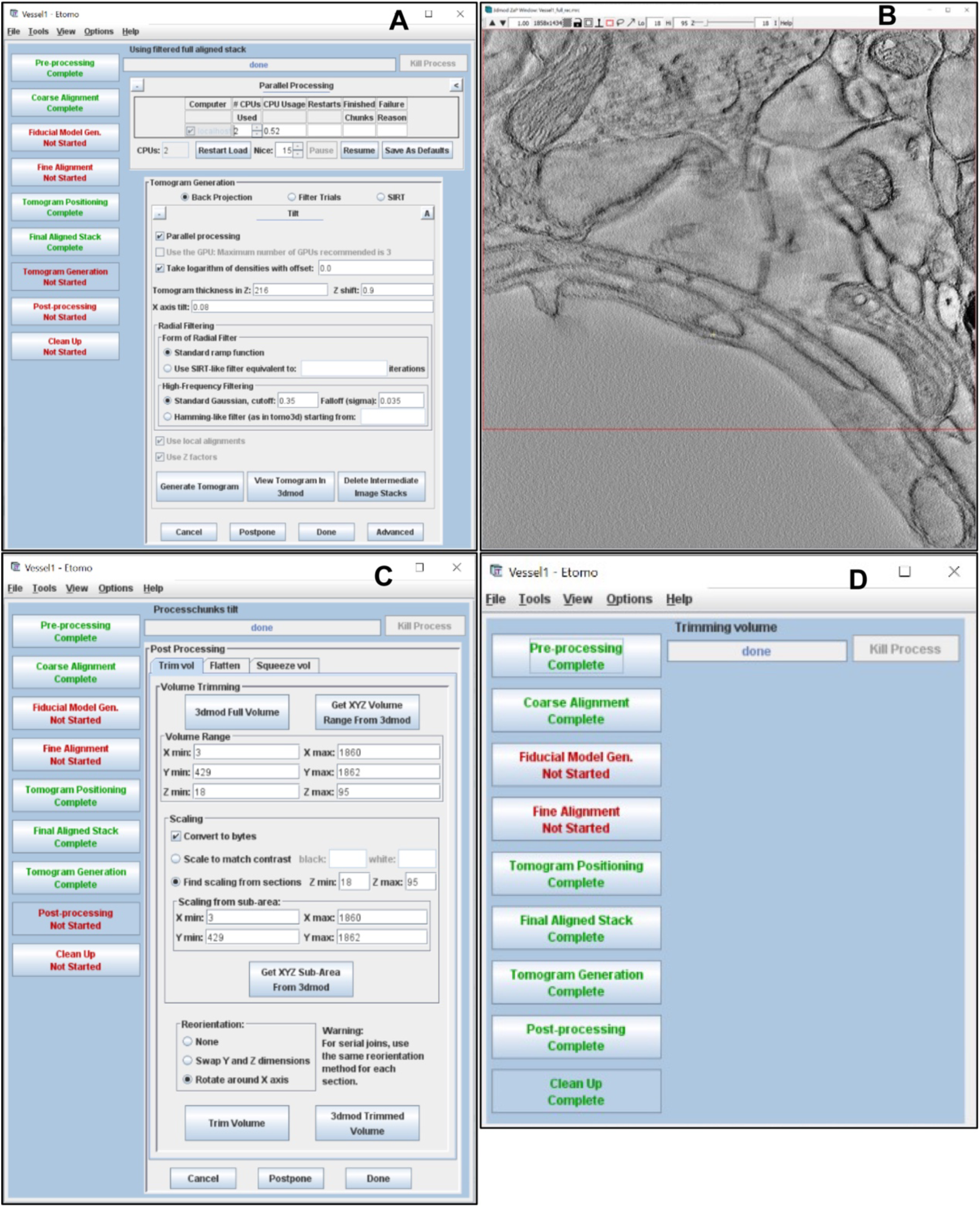
Tomogram generation. A) Application of weighted back projection for tomogram generation. B) Trimming process of generated tomogram. Red rectangle delineates the cropped area. C) Adjustment of tomogram coordinates after trimming. D) Final window of tomogram generation.

After defining the optimal Z range and region of interest, navigate back to the main Etomo window without closing previous window and press “Get XYZ Volume Range From 3dmod”. The software will adjust the tomogram parameters according to the trimming step. Afterwards, select “Get XYZ Sub-Area From 3dmod” and press “Trim Volume”. Finally, press “Done” to generate a trimmed 3D reconstructed volume (**Figure 4C**). The final step of tomogram generation consists of cleaning up the intermediate files created during the process (**Figure 4D**). The generated tomogram will be saved automatically as “*name_rec.mrc*” in the same directory containing the raw image stack *.st* file and will be used for further segmentation step.

### 5.4 Tomogram segmentation

Tomogram segmentation is performed manually using 3dmod software^64^. As AI algorithms designed to assist with segmentation of brain endothelial cells are still missing, the manual method remains to option of choice for greater accuracy. Training data requirements and the inherent complexity of biological structures make the development of AI-based segmentation methods highly challenging.

To initiate tomogram segmentation, the user should launch 3dmod software by entering the command “3dmod” in the command line. A 3dmod startup window will appear; select the “*name_rec.mrc*” file containing the reconstructed 3D volume. By pressing “OK,” a new 3dmod ZaP window will appear, displaying the reconstructed volume with the Z-axis range on top. Prior to starting the tomogram segmentation, ensure that the mouse first, second, and third buttons are set to “Right, Left, Middle**”. To do this, press “3dmod”, situated on the left upper corner of the menu, then select “Preferences → Done”. If the 3D volume appears noisy, this can be resolved through the Fourier filter by selecting “Edit → Image → Process.” Reducing “High-frequency cutoff” and “High-frequency falloff” parameters will minimize image noise. However, this will also affect resolution, which might induce bias regarding fine ultrastructures. Therefore, it is crucial to maintain a balance between these parameters to preserve faithfulness of the reconstruction. Once the mouse settings and image quality are set appropriately, ensure the “Model” mode is selected in the main 3dmod window. Before starting segmentation, press “Image → Model View”. Create a new object by pressing “Edit” located on the menu bar and selecting “Object → New”. After attributing the name of the structure of interest (*i.e.*, endothelial cell 1), select the “Closed” object type to facilitate further contour interpolation. Of note, the number of created objects should correspond to the number of the ultrastructures of interest. After setting all the parameters, press the “Special” button located on the menu bar and select “Drawing tools → Normal” to draw the first contour of the first structure of interest. We recommend drawing the contours of endothelial cells first and subsequently proceeding to the segmentation of smaller ultrastructures (*i.e*., transcytotic vesicles, ER). To zoom-in onto the specific region of the image, or zoom out, press a combination “CTRL + mouse wheel” for WindowsOS or “command + mouse wheel” for MacOS to ensure precise contour drawing. Additionally, zooming in on the contour may exhibit irregular shapes. To address this, select the “Sculpt” drawing tool to smooth or correct all the irregularities. Manually drawing every contour on each slice is a time-demanding process, especially for big datasets. Interpolation helps to minimize the time of the segmentation process by automatically generating the contours across the gap between manually drawn contours of non-adjacent slices. To use this tool, navigate to the menu bar of the 3dmod ZaP Window and press “Special → Interpolator”. A new window will appear, offering the following interpolation methods: “Linear”, “Spherical”, “Smooth” or “Smooth pts” which refers to smooth points. The choice of interpolation type depends on the contour shape. Briefly, “Linear” interpolation is used for straight lines, “Spherical” interpolation is dedicated to the generation of spheres, and “Smooth” type is used for curvilinear lines. For endothelial cells, we recommend using either “Smooth” or “Linear” interpolation, depending on the structural complexity. For cytoplasmic vesicles, the “Spherical” type is optimal. Furthermore, to ensure contour accuracy and continuity between slices, we recommend using an interpolator tool after manually drawing contours on every fifth slice. Detailed information regarding the use of the interpolators is provided by plugin help, located in the upper right corner.

The newly generated contours can be visualized by navigating to “Image → Model View” or by pressing the “V” button. Once the contour ensemble of each object appears consistent, it can be meshed into the homogenous surface by navigating to the “Meshing” panel. Each object can be meshed separately by selecting the object and pressing the “Mesh One” button, or all the objects can be meshed at once by pressing “Mesh All”. To verify the consistency of meshed surfaces, rotate the model by holding the middle mouse button. The contours can be smoothed by checking “Surface”, increasing the “Z inc” value, and then completing the meshing process. Alternatively, the contours can be refined by using the “Sculpt” drawing tool in a main 3dmod ZaP Window, as described above. To reset the model orientation, press the “T” button.

In addition, the model can be complemented by superposing it onto the tomogram by navigating to the menu bar, selecting “Edit → Image", then pressing the “View Z image” option, or pressing the “Shift + Z” combination. Contrast and transparency can be adjusted automatically by selecting “Use 3dmod Black/White”.

Once the entire rendered 3D model appears consistent and correctly oriented, it can be acquired by selecting ‘HotKeys” in the menu bar and press “Make TIFF snapshot of window” or “Make JPEG snapshot of window” or “Make PNG snapshot of window”. To display scalebar, press “View → Scale Bar…”. The acquired snapshots will be saved automatically in the following directory “Computer → Macintosh HD → Users → username folder” for MacOS. For WindowsOS, prior to acquire snapshot, set the snapshot directory by selecting “File” in the main 3dmod window and pressing “Set Snap Dir..”. Furthermore, the model can also be acquired for movie generation. Firstly, select the “Model” mode in the main 3dmod window. Then, select the 3dmod Model View window and “File → Movie/Montage” or press the “M” button. A new “MV Movies” will display the parameters for movie creation, such as rotations at different axes, translations, and navigation through slices. After selecting the desired parameters, adjust the “# of movie frames” and the “FPS”. For our movie, we set “70” for the “# of movie frames” and “15” for “FPS”. After setting all the parameters, the movie can be previewed before being created by pressing “Make”. Once all the settings are satisfying, check “Write files” and press “Make”. The sequence of snapshots will be saved automatically into the same directory as the snapshots.

We use Fiji software (NIH) to compile the sequence of snapshots into a movie. Launch the software and press “File → Import → Image Sequence…” then select the folder containing the acquired sequence of snapshots and press “OK”. A new Fiji window displaying compiled snapshots will appear, and the movie is ready to be saved by pressing “File → Save As… → AVI…”. Choose the desired compression and set the frame rate corresponding to the “FPS” value chosen in the “MV Movies” window. After selecting “OK”, choose the directory where the movie file should be saved.

## Conclusion

Brain endothelial cells form a unique selective and protective barrier, known as the BBB, limiting the entrance of blood-borne molecules into the brain. In pathological conditions, the BBB is disrupted, resulting in neuronal damage. Understanding the fine changes underlying BBB disruption requires advanced imaging tools. TEM is a powerful tool that allows ultrastructural investigation of ECs at the BBB. While conventional TEM is limited to 2D single-layer snapshots of thin specimens, ET emerges as a method to overcome this major limitation, offering detailed views of specimen in 3D at the ultrastructural level. Here, here present the key concepts of TEM and RT-ET, and include a detailed protocol to guiding readers from sample preparation to 3D reconstruction of brain endothelial cells, with a focus on inter-endothelial junctions. Doing so, this method paper aims to facilitate understanding of the ET process, thereby expanding research tools to investigate microvascular biology in health and disease context at nanoscale resolution.

